# Conformational stability and dynamics in crystals recapitulate protein behaviour in solution

**DOI:** 10.1101/2020.04.27.063909

**Authors:** BM Sala, T Le Marchand, G Pintacuda, C Camilloni, A Natalello, S Ricagno

**Affiliations:** Royal Insitute of technology; Centre de RMN à Très Hauts Champs, Institut de Sciences Analytiques; Università degli studi di Milano; University of Milano-Bicocca; University of Milan

**Keywords:** Molecular Dynamics, FTIR spectroscopy, protein stability

## Abstract

A growing body of evidences has established that in many cases proteins may preserve most of their function and flexibility in a crystalline environment, and several techniques are today capable to detect transiently-populated states of macromolecules in tightly packed lattices. Intriguingly, in the case of amyloidogenic precursors, the presence of these conformations (hidden to conventional crystallographic studies) can be correlated to the pathological fate of the native fold.

It remains unclear, however, to which extent these minor conformations reflect the protein behaviour that is more commonly studied in solution. Here, we address this question by investigating some biophysical properties of a prototypical amyloidogenic system, β2-microglobulin (β2m) in solution and in microcrystalline state.

By combining NMR chemical shifts with Molecular Dynamics (MD) simulations, we confirmed that conformational dynamics of β2m native state in the crystal lattice is in keeping with what observed in solution.

A comparative study of protein stability in solution and *in crystallo* is then carried out, monitoring the change in protein secondary structure at increasing temperature by Fourier transform infrared (FTIR) spectroscopy. The increased structural order of the crystalline state contributes to provide better resolved spectral components compared to those collected in solution and crucially, the crystalline samples display thermal stabilities in good agreement with the trend observed in solution.

Overall, this work shows that protein stability and occurrence of pathological hidden states in crystals parallel their solution counterpart, confirming the interest of crystals as a platform for the biophysical characterisation of processes such as unfolding and aggregation.

## Introduction

The crystallisation of proteins has allowed one of the major revolutions in the understanding of macromolecules: to date X-ray crystallography has elucidated more than 160,000 protein structures (www.rcsb.org) and it is the reference method for structural biology. Protein molecules are rather loosely packed in crystal and the average solvent content is 50% (1). Crucially this allows enough conformational freedom so that many enzymes are active in crystalline form (2, 3) and many processes which entail internal rearrangements and minor conformational changes may be monitored *in crystallo* by serial synchrotron crystallography and time-resolved crystallography (4–6). However, although in crystal structures poor electron density and high B-factors indicate that protein molecules may retain rather dynamic and flexible conformations in crystals, the common view is that the study of protein crystals is limited to the determination of static structures.

Solid-state NMR offers the unique possibility to evaluate protein dynamics in crystals by using spin relaxation techniques to probe motions with site and time specificity (7–10). Since the first pioneering studies which quantified molecular motions in crystals, critical evaluation of the equivalence between such dynamics and those happening in solution have been performed, providing pictures that differ according to the motional processes under investigation. Fast motions typically happening in the pico-to nanoseconds regime, have been demonstrated to be similar in solution and in crystals (11–13). Importantly, solid-state NMR is very sensitive to slower collective motions, and has revealed the presence of excited states in exchange with the main conformational state of the molecule at a rate of micro-to milliseconds in the crystal lattice (14). The possibility to detect such transient conformations is of tremendous interest as they are associated to processes such as molecular transport, allosteric regulation, folding/unfolding and aggregation. The amplitude and timescale of conformational exchange processes may be affected by crystal packing, nevertheless they have been shown to directly report on the corresponding motions happening in solution (15).

While many of these data have been performed on model crystalline samples, we recently investigated the molecular bases of D76N β2-microglobulin (β2m) pathologic amyloid propensity by studying protein dynamics in crystals (16). Combining solid-state NMR and replica-averaged metadynamics ensemble simulation, we showed that the presence of ‘hidden’ conformations and increased dynamics in crystals correlates with amyloid formation (16).

The fact that biophysical properties of a protein are comparable in crystallo and in solution, however, appears as counter-intuitive to part of the community, in particular in the challenging context of misfolding and aggregation. Specifically, native states and pathological conformations may be connected by large-amplitude conformational changes, which are likely affected by intermolecular contacts, even in a largely hydrated lattice. It remains therefore unclear to which extent the minor conformations detected in crystals reflect the protein behaviour that is more commonly studied in solution. In the attempt to clarify this aspect, we perform here a comparative study of protein dynamics and fold stability of the prototypical system β2m in the crystalline state and in solution.

β2m is the light chain of Major Histocompatibility Class I complex (MHC-I), well known as an aggregation-prone protein associated with two different amyloid-related pathologies (17). Wild type (wt) β2m is the causing agent of an amyloidosis affecting patients undergoing long-term haemodialysis, condition referred as dialysis-related amyloidosis (DRA) (18). The D76N genetic variant, instead, is responsible for a familial systemic amyloidosis (19), and shows increased aggregation propensity as compared to the wt protein (19, 20). The remarkable properties of D76N β2m can be related to the perturbation of a crucial network of weak intramolecular interaction resulting in increased dynamics, lower protein stability and increased amyloidogenicity for a native-like excited state (16). Conversely, an artificial mutant of the evolutionary conserved Trp60 displays a much lower aggregation propensity (21). Given the relevance of structural and biophysical properties for amyloid aggregation, such three β2m variants have been studied in details. Noteworthy, all three β2m variants crystallise under identical chemical-physical conditions and form the intermolecular interactions in identical crystal packing, avoiding interpretation biases of the solid-state data.

The conformational exchange processes of D76N and wt β2m are simulated based on the inclusion of NMR chemical shifts determined either from crystalline or solution samples. Moreover, protein stability was assessed for the set of the three mutations (D76N, wt and the highly stable W60G), using Fourier transform infrared (FTIR) spectroscopy, a biophysical technique which is amenable for monitoring both solid and liquid samples by following the changes in secondary structure. FTIR spectroscopy has been widely adopted for the analysis of protein samples in different conditions, resulting a well suitable technique to compare the secondary structures of proteins both in solution and in crystals as well as in intact cells, tissues and entire organisms (22–28). In particular, a recent work published by our group showed the feasibility of collecting FTIR spectra on single protein crystals and highlighted the high quality of the FTIR spectra collected on β2m crystallin samples (29). Our results spot differences between solution and crystals, consistent with a systematic increase in protein stability in the crystal lattice. At the same time, however, they explicitly indicate a persistence of dynamical properties between the two environments, with conserved relative conformational stabilities across a set of β2m mutants. This study reinforces a picture where crystals contain a wealth of information about intrinsic properties of proteins as dynamics and stability and that deep insights into phenomena such as unfolding and aggregation can be inferred from native conformations in crystalline samples.

## Statement of Significance

Proteins preserve their function and flexibility in a crystalline environment, but it is still unclear whether their properties in crystallo reflect protein behaviour in solution. To address this question, in this work we have combined NMR chemical shifts with Molecular Dynamics (MD) simulations and conformational ensembles of protein molecules in crystallo and in solution were calculated. Moreover, protein stability was compared in solution and *in crystallo* by Fourier transform infrared (FTIR) spectroscopy. Here we show that crystalline protein samples can provide valuable information beyond crystal structures. This work highlights the potential value of crystals as a platform for the biophysical characterisation of proteins giving possibly access to hidden conformational states related to pathological conditions.

## Materials and Methods

### β_2_-microglobulin expression and purification

All β_2_m variants were expressed in BL21 (DE3) pLysS *E. coli* strain as inclusion bodies. The proteins were extracted and purified following previously published protocols (21). For NMR studies, ^13^C, ^15^N and ^2^H enrichment was obtained as described in(16).

### Solution NMR of D76N β2m

For NMR measurements, a solution of triply-labelled ^1^H^N^,^2^H,^13^C,^15^N,-D76N β_2_m was prepared at a concentration of 100 μM in a 70 mM sodium phosphate buffer, at pH 7.6. NMR experiments were recorded at 293 K on Bruker Avance 600 Spectrometer equipped with a TCI cryoprobe operating at a ^1^H Larmor frequency of 600.6 MHz. NMR spectra were referenced to the DSS for ^1^H and indirectly for ^13^C and ^15^N, accounting for the isotope effect due to the deuteration of the protein, as recommended by IUPAC. Site-specific resonance assignment has been determined on the basis of a 2D ^1^H,^15^N-HSQC and 3D HNCO, HNCA, and HNCACB experiments. Manual assignment was performed with the program CARA (http://cara.nmr.ch). Chemical shifts are available under BMRB entry 50223.

### Molecular dynamics simulations

Simulations data for wt β2m and D76N β2m in crystals were taken from (16), simulations data for wt β2m in solution were taken from (30), where all simulations were run following comparable procedures. D76N β2m simulation in solution were newly performed using the corresponding NMR chemical shifts, following the protocol briefly reported in the following. Simulations were carried out using GROMACS and PLUMED with the ISDB module. The system was described using the Amber03W force field in explicit TIP4P05 water at 298 K. The starting conformation was taken from PDB 4FXL. The structure was protonated and solvated with ~8200 water molecules in a dodecahedron box of ~260 nm^3^ of volume. The metadynamics metainference (M&M) protocol was applied using chemical shifts and a global outlier model for the noise as previously described. Thirty replicas of the system were simulated in parallel with a restraint applied on the weighted average value of the back-calculated NMR chemical shifts with a force constant determined on the fly by M&M.

All replicas were biased by Parallel Bias Metadynamics along the following four collective variables (CVs): the antiparallel beta content (the “anti-β” CV), the AlphaBeta CV defined over all the chi-1 angles for the hydrophobic side-chains (the “AB” CV), the AlphaBeta CV defined over all the chi-1 angles for the surface exposed side-chains (the “ABsurf” CV), and the AlphaBeta CV defined over all the phi and psi backbone dihedral angles of the protein (the “bbAB” CV). Definition of the CVs are available in the PLUMED manual. Gaussians deposition was performed with σ values automatically determined by averaging the CV fluctuations over 2000 steps and setting a minimum value of 0.1, 0.12, 0.12, and 0.12, for anti-β, AB, ABsurf, and bbAB, respectively; an initial energy deposition rate of 2.5 kJ/mol/ps and a bias-factor of 20. Furthermore, to limit the extent of accessible space along each collective variable and correctly treat the problem of the borders, intervals were set to 12–30, 10–40, 0–33, and 10–42 for the four CVs, respectively. Each replica has been run for a nominal time of 350 ns.

The sampling of the 30 replicas was combined using a simple reweighting scheme based on the final metadynamics bias *B* where the weight *w* of a conformation *X* is given by *w=exp(+B(X)/k_B_T)*, with *k_B_* the Boltzmann constant and *T* the temperature, consistently with the quasi static behaviour at convergence of well-tempered metadynamics. The convergence of the simulations by block analysis, including error estimates, is shown in **Figure** S4. All the data and PLUMED input files required to reproduce the results reported in this paper are available on PLUMED-NEST (www.plumed-nest.org), the public repository of the PLUMED consortium as **plumID :20.001**.

### Crystallization and sample preparation for Fourier transform infrared (FTIR) spectroscopy

Lyophilized wt, W60G and D76N β2m variants were solubilized in ddH2O at a concentration of 8.5 mg/ml. All mutants were crystallized with sitting drops technique at 20°C. 160 μl of protein solution was mixed in a bridge with the same amount of 0.1 M MES pH 6, 27% PEG 4K, 15% glycerol, and placed against 0.1 M MES pH 6, 30% PEG 4K, 15% glycerol as reservoir solution. In few days, needle-like crystals were grown. All three variants not only crystallise under the same conditions but also crystals have the same space group and crystal packing (19, 21). Thus, all underlying intermolecular interactions in the crystals are identical in the three β2m variants. This makes data obtained from crystals comparable.

Before FTIR measurement the crystallization drop was collected and centrifuged at 2000 rcf for two minutes. Once removed the supernatant, the crystal pellet was washed and centrifuged three times with 0.1 M MES pH 6, 27% PEG 4K, 15% glycerol, all component solubilized in deuterated water to achieve the full D to H exchange of all exchangeable H. Crystals were then incubated overnight in 350 μl of the same deuterated buffer and washed once again before measurement.

### FTIR of β2m crystals

For the FTIR analysis, crystals were resuspended in 20 μl of deuterated buffer (0.1 M MES pH 6, 27% PEG 4K, 15% glycerol) and then transferred in a temperature-controlled transmission cell with two BaF_2_ windows separated by a 100 μm Teflon spacer.

FTIR spectra were collected at room temperature (~25°C), in transmission mode, using a Varian 670-IR spectrometer (Varian Australia, Mulgrave, Australia), equipped with a nitrogen-cooled mercury cadmium telluride detector, under accurate dry air purging. Spectra were collected under the following conditions: 2 cm^−1^ resolution, 25 kHz scan speed, 1000 scan coaddition, and triangular apodization (29).

Thermal stability experiments were carried out heating the sample from room temperature to 100°C at a rate of 0.4 °C/min and 1 °C/min, collecting each transmission spectrum every ⁓3.4 °C and ⁓4.3 °C respectively. The FTIR spectra were obtained after subtraction of solvent absorption, collected under the same conditions. For a better comparison, the absorption spectra were normalized at the same Amide I band area before spectral smoothing (25 points) and second derivative calculation. Collection and analysis of the FTIR spectra was performed by the Resolutions-Pro software (Varian Australia, Mulgrave, Australia)

### FTIR of β2m variants in solution

Lyophilized β2m was dissolved in deuterated buffer 50 mM Sodium Phosphate pH 7.4, 100 mM NaCl, at a final concentration of 2.5 mg/ml. 20 μl of the sample were transferred in a temperature-controlled transmission cell with two BaF_2_ windows separated by a 100 μm Teflon spacer. Thermal stability experiments and spectral analyses were performed as described above for the crystalline pellet (29).

## Results

### NMR-restrained molecular dynamics: comparison between conformational ensembles in solution and in crystals

Conformational dynamics of β2m in crystals and in solution were analysed by combining NMR chemical shifts and MD simulations in the theoretical framework of Metainference. In previous works we have characterised the conformational ensembles for wt and D76N β2m in crystal and for wt β2m in solution (16, 31). Here, we complement the analysis by collecting NMR chemical shifts in solution for D76N β2m (**Fig. S1**), and deriving the corresponding conformational ensembles.

The ensembles derived from the MD simulations were first analysed in terms of the ‘Anti-beta’ parameter, proportional to the number of beta-sheet elements and the configuration of the solvent-exposed sidechains (‘Sidechain’), which have been shown to be important descriptors of the aggregation potential (16). From the corresponding Free Energy Surfaces (FES), which are reported in **Figure** 1, it appears clear that the overall description is highly conserved between solution and the solid state. The data show the presence of a higher-energy conformational ensemble in D76N β2m in solution (M_1_*), in line with what has been observed for the three other cases. In qualitative agreement with our previous findings in crystals, the beta-sheet structure destabilisation in the excited state M_1_* is larger in D76N as compared to wt β2m and the relative population of this state is lowered by the mutation. Both ground and excited states of the two proteins are more structured (as monitored by the higher ‘Anti-beta’ parameter) in the crystals than in solution, mirroring what is typically observed between structures obtained by X-ray crystallography and solution NMR of the same protein.

**Figure 1.**
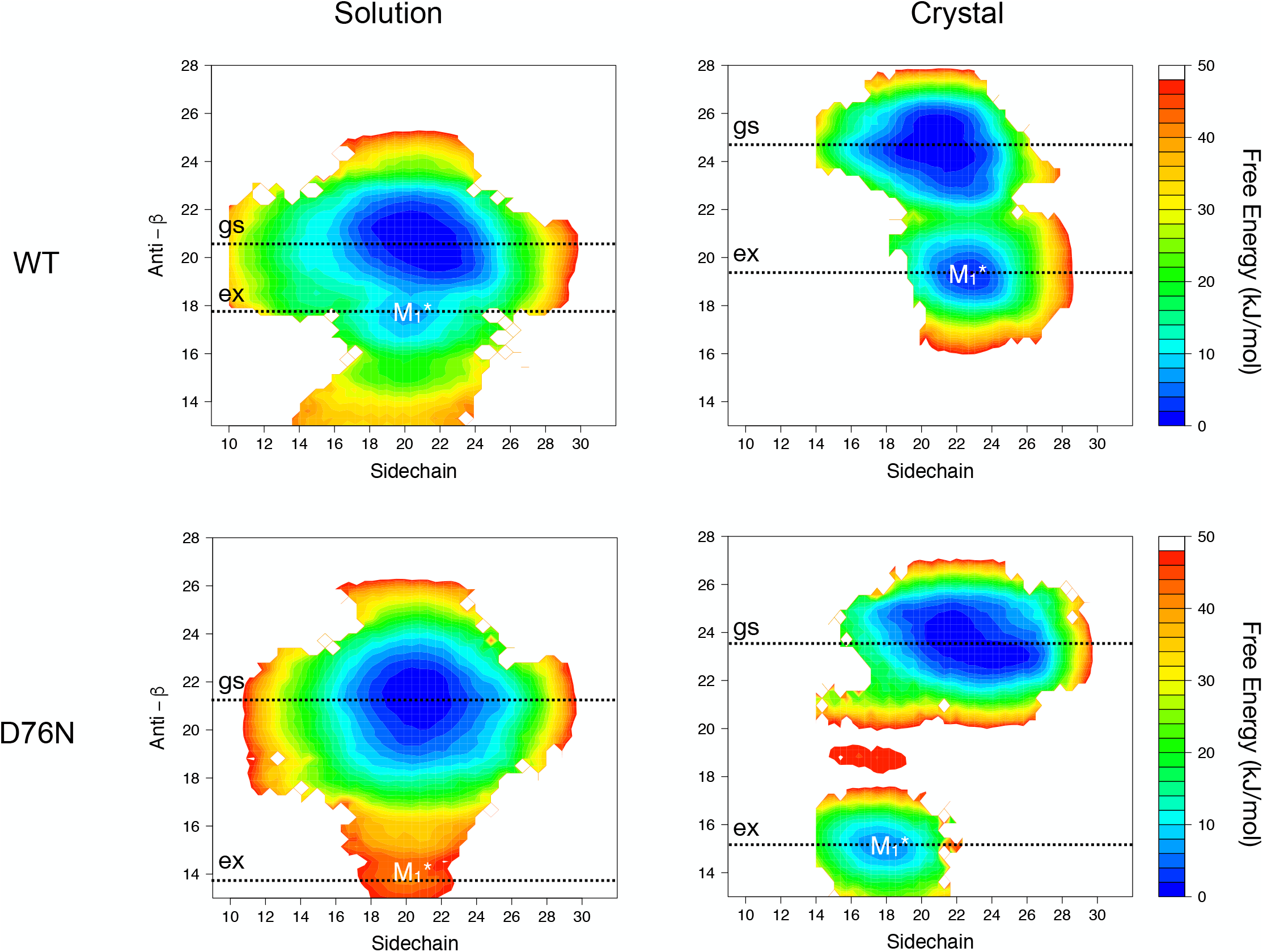
Conformational ensembles for native β2m variants in solution and *in crystallo*. Free Energy Surfaces (FES) in kJ/mol for the conformational ensembles of wt β2m and D76N β2m using solution or solid-state NMR chemical shifts as experimental restraints. The FES are shown as a function of the Anti-b RMSD, which is proportional to the amount of beta structure, and of the Sidechain CV, which reports about the overall configuration of the solvent exposed side-chains. FES for wt β2m in solution is taken from (30), the FES for WT- and D76N-in the solid state are from (16), while the FES for D76N β2m in solution is reported here for the first time. The dotted lines indicate the position of the minima in the FES.

We subsequently investigated a second key parameter for β2m fold stability, the solvent accessibility of residue W95, a crucial residue for β2m hydrophobic core (21). The proximity of W95 to the site of the D76N mutation may result in an increased accessibility to the solvent in the D76N variant as compared to wt β2m. The analysis of the X-ray structures did not support this hypothesis (19) as well as the average Solvent Accessible Surface Area (SASA) calculated over the ensembles did not show a clear difference between wt and D76N indicating W95 as fully buried. Here we re-analysed our former and newly determined conformational ensembles to check for the presence of low-populated states that may show an increase W95 accessibility. In **Figure** 2 we report the free-energy surfaces for the four conformational ensembles as a function of the beta content (as in **Figure** 1) and of W95 SASA. Crucially, a second minor high-energy state highlighted in **Figure** 2 by a vertical line, distinct from the one previously discussed, is present in D76N and it is characterised by rather high beta-content and by a highly solvent exposed W95 (M_2_*). This state is not present in the wt β2m projections. The high-energy state M_1_* was suggested to be associated with protein misfolding and consequently with aggregation (16, 32, 33). Conversely, given the pivotal role of W95 in the stability of β2m buried hydrophobic core (21, 34), one may speculate that excited state M_2_*, having a highly solvent exposed W95 residue, may be associated with protein folding-unfolding and consequently with thermodynamic stability.

**Figure 2.**
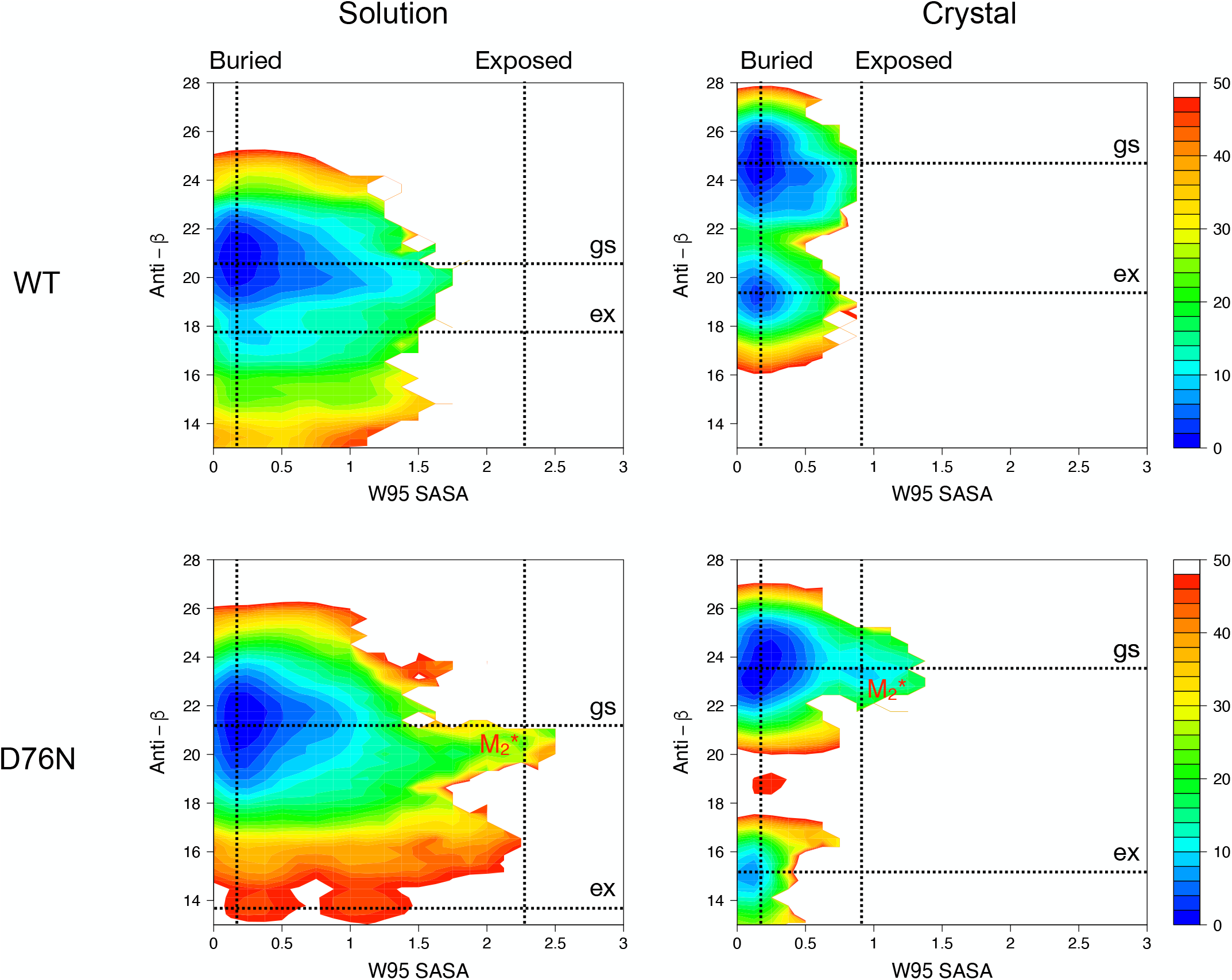
W95 role in β2m’s conformational ensembles. Free Energy Surfaces (FES) in kJ/mol for the conformational ensembles of wt β2m and D76N β2m using solution or solid-state NMR chemical shifts as experimental restraints. The FES are shown as a function of the Anti-b RMSD, which is proportional to the amount of beta structure, and of the Solvent Accessible Surface Area (SASA) in nm^2^ of the W95 residue. The dotted lines indicate the position of the minima. This representation shows how both the previously identified ground and excited state are characterised by a buried W95 but that in the case of D76N there are minima associated with a solvent exposed W95 compatible with the lower protein stability.

### FTIR spectroscopy: comparison between β2m stability in solution and in the crystalline state

Above, we examined the molecular determinants of the unfolding and amyloidogenic properties of β2m in solution and in crystals and found quantitative conservation between the two states. We then focussed on if and how the crystal packing affects β2m stability. In order to assess whether such biophysical property may be monitored on crystalline samples, a comparative secondary structure characterisation of crystalline and soluble β2m variants was carried out by FTIR spectroscopy. In order to have a more representative set of protein stability values, the highly stable W60G β2m mutant (21) was added to the two β2m variants analysed above.

First FTIR experiments have been performed (see below) in the crystallization solution to better compare the resulting data with the ones obtained from the crystalline state. The spectra collected for the soluble native proteins in crystallization conditions display the same spectral features of those collected in phosphate buffer (29), indicating comparable secondary structures (**Figure** 3A). However, under such conditions, proteins tend to precipitate leading to low quality non-reproducible measurements (data not shown). Thus, high quality spectra and temperature ramps have been carried out in deuterated phosphate buffer for the soluble β2m variants while proteins in the crystalline state have been studied in deuterated crystallization solution, as described in Materials and Methods section.

**Figure 3.**
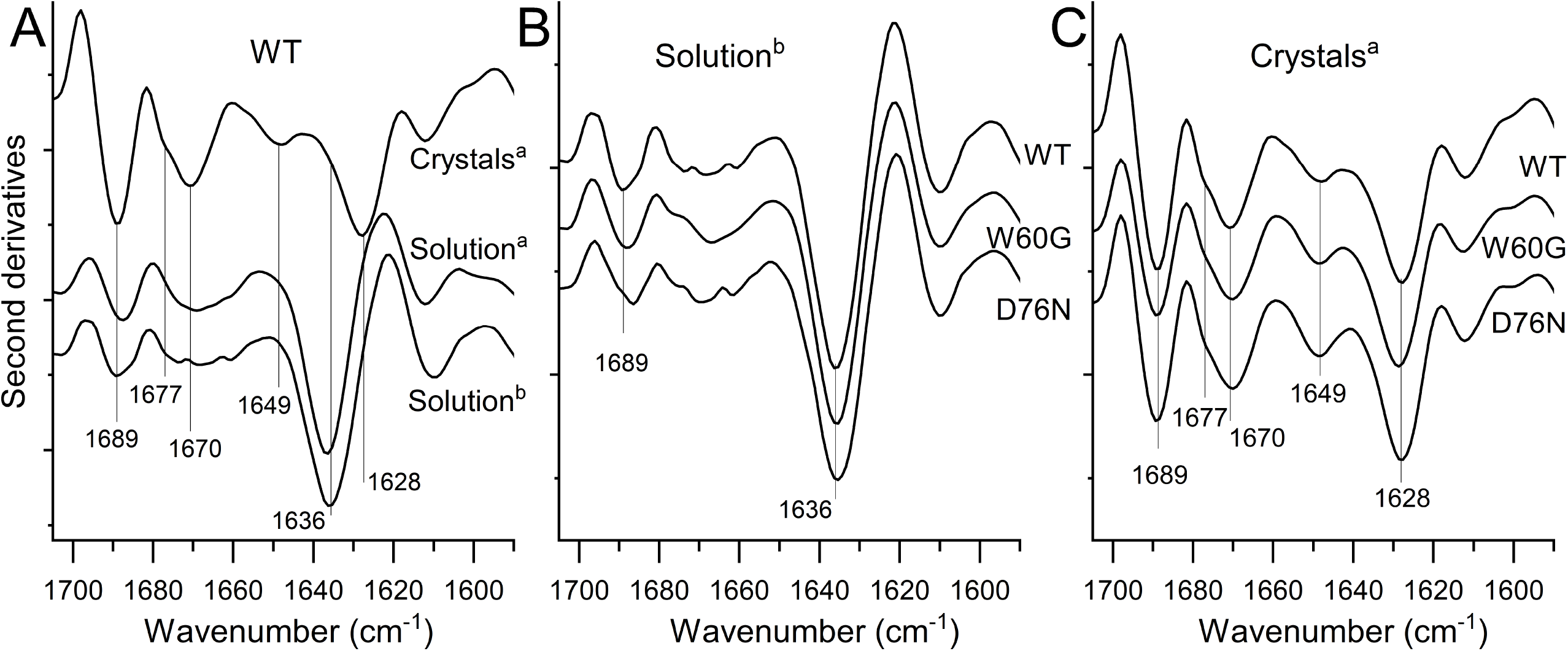
FTIR spectroscopy of wt, D76N and W60G β2m variants in solution and in the crystalline state. **(A)** The second derivatives of the absorption spectra of wt β2m in solution compared with that of wt protein in the crystalline state. **(B)** Comparison of the second derivative spectra of wt, D76N and W60G β2m in deuterated phosphate solution. **(C)** Comparison of the second derivative spectra of wt, D76N and W60G β2m crystals in deuterated crystallization solution. The peak positions of the main Amide I components are indicated. ^a^ Samples in deuterated crystallization solution; ^b^ Sample in deuterated phosphate solution.

**Figure** 3A shows the second derivatives in the Amide I region of the FTIR absorption spectra collected at room temperature for the wt β2m protein in solution and in the crystalline state. The Amide I band is due to the C=O stretching mode of the peptide bond and it is particularly sensitive to the protein secondary structures, including the intermolecular β-sheet structures in protein aggregates (26, 27, 29, 35–37). The second derivative spectrum of crystalline wt β2m displays sharper Amide I peaks suggesting more rigid protein molecules compared to what observed in the solution spectra (**Figure** 3A). The peak due to the native antiparallel β-sheets is downshifted from ~1636 cm^−1^ in solution to ~1628 cm^−1^ in the crystals, likely reflecting a stronger hydrogen bonding in the β-sheets of the protein related to the crystal packing (29, 35). Due to the increased structural order of the crystalline state, new and better-resolved bands are detectable in the crystalline sample (29). In fact, the turn absorption band at ~1670 cm^−1^ is narrowed in wt β2m crystalline sample compared to solution. Moreover, two closed components at ~1689 and the new at ~1677 cm^−1^ have been assigned to intramolecular β-sheet structures and/or turns, while the new band at ~1649 cm^−1^ in the crystal has been related to loop regions(22, 29, 35). These results are in agreement with a previous study on β2m variants where FTIR micro-spectroscopy was used to characterise single β2m crystals transferred in D_2_O (29). In the present study, instead, bulk crystalline samples were transferred in deuterated crystallization buffer and measured by a temperature-controlled IR cell after over-night incubation at 4°C (36).

Spectra of the D76N and W60G variants were also collected both in solution and in crystalline state and compared to the analyses carried out on WT β2m (**Figure** 3B and C). As mentioned above for the wt protein, the same relevant differences are observed between the spectra measured in solution and in crystals. Indeed, better resolved components are well detectable together with the downshift of the main native β-sheet band, in agreement with our previous study (29). Neither the comparison of spectra taken from crystalline samples nor the one from samples in solution detect significant differences between the three variants.

### Temperature ramps of the WT, D76N and W60G variants in solution and in the crystalline state

The thermal stability of the β2m variants in solution and in the crystalline states was studied by FTIR spectroscopy heating the samples from room temperature (RT) to 100°C. In both crystalline and solution states, the three variants show that the loss of native β-sheet structures is followed by protein aggregation, as demonstrated from the appearance of new bands at ~1619 and 1684 cm^−1^ (**Figure** 4A-I and **Figure** S2), related to the formation of intermolecular β-sheet structures, as previously reported (29).

**Figure 4.**
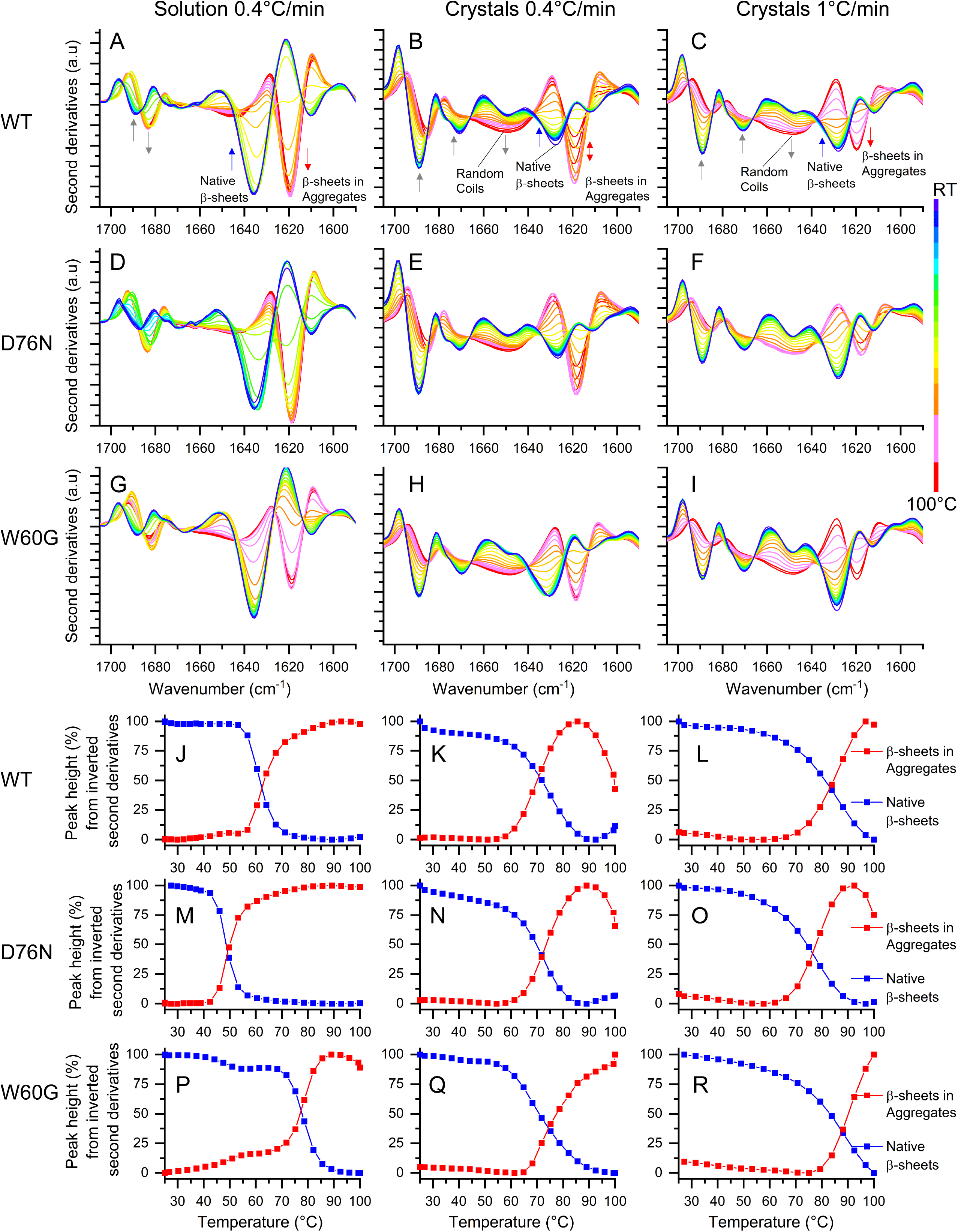
Temperature dependence of FTIR spectra of the β2m variants in solution and in crystals. **(A-I)** Second derivatives of the absorption spectra collected at increasing temperature. The β2m variants were analysed in solution in deuterated phosphate buffer (left panels) and in the crystalline state (middle and right panels) in crystallization conditions. The heating rates were indicated. The assignment to the protein secondary structures of the main Amide I components is reported in the upper panels. The arrows point to increasing temperatures. (J-R) Temperature dependence of the native β-sheets (blue) and of β-sheets in protein aggregates (red) evaluated from the peak intensities in the second derivative spectra of panels A-I, respectively.

Taking the D76N variant as a representative case for the spectral changes observed in solution, the native β-sheet component at ~1636 cm^−1^ was found to be stable up to ~42°C and to rapidly decrease above this temperature (**Figure** 4D, 4M). The loss of the native β-sheet structures was accompanied by the raising of the ~1619 cm^−1^ and 1684 cm^−1^ peaks (**Figure** 4D, 4M). As shown in **Figure** 4A and 4G, thermal treatments induced similar spectral changes for the wt and W60G β2m in solution compared to D76N solution. The temperature dependencies of the native β-sheet peak height and of that of the β-sheets in protein aggregates for the three variants in solution are shown in **Figure** 4 J M and P. The calculated mid-point temperatures (*T*_*mp*_) inferred from the thermal denaturation curves of the native structures are reported in **Figure** 5B. The *T*_*mp*_ values indicate the following stability trend: W60G (~79.8°C) > wt (~ 63.6°C) > D76N (~ 51.5 °C), which is in fully agreement with previous results obtained by different spectroscopic approaches (29, 38).

**Figure 5.**
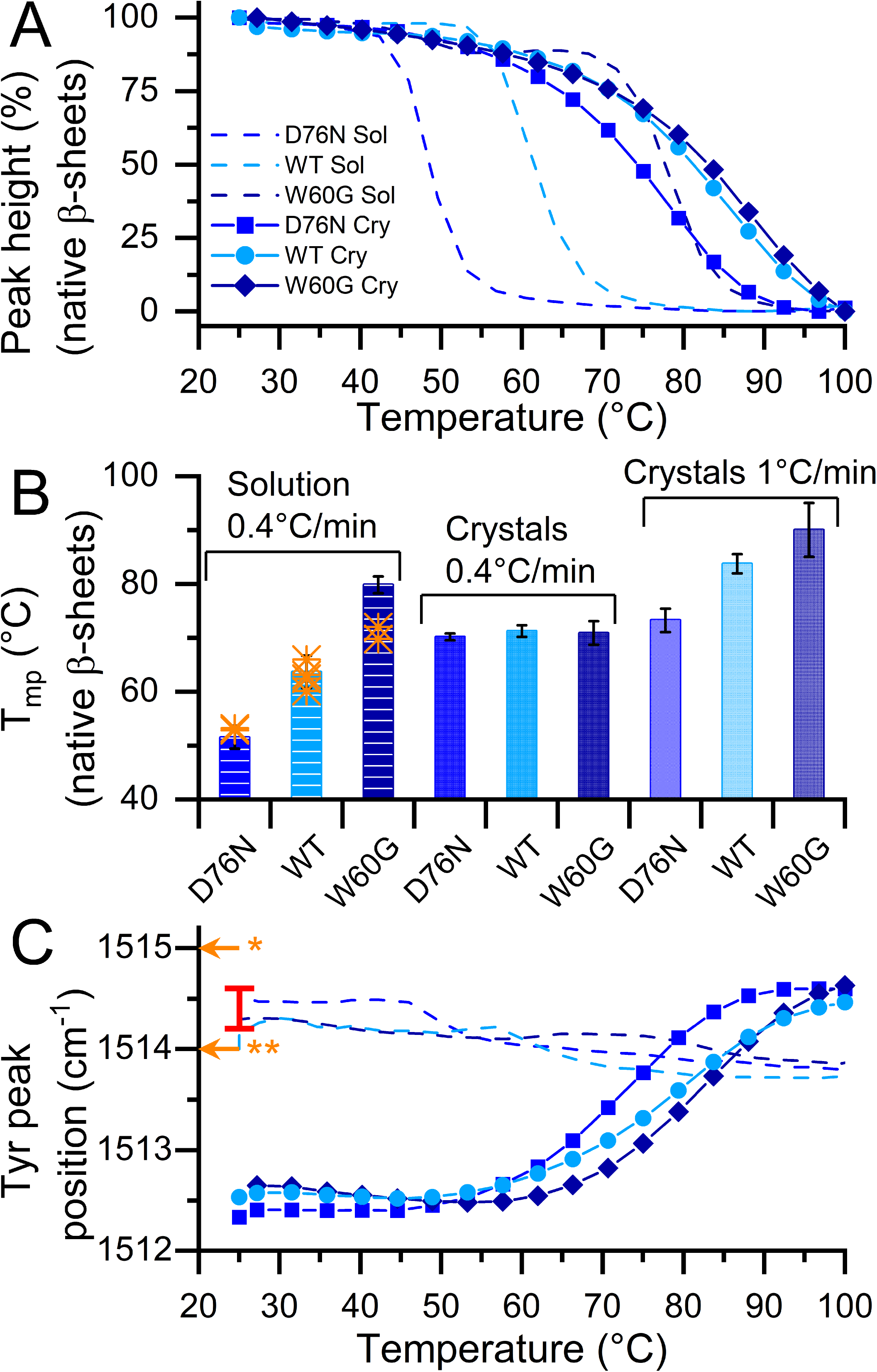
Thermal stability of β2m variants assessed by FTIR spectroscopy. **(A)** The temperature dependence of the native β-sheet peak height of β2m variants in solution and in the crystalline state. Proteins in solution were heated at 0.4°C/min and the intensities of the ~1636 cm^−1^ peak, taken from second derivative spectra, were reported. For protein crystals, samples were heated at 1°C/min and the intensities of the ~1628 cm^−1^ peak, taken from second derivative spectra, were reported. **(B)** Calculated mid-point transition for the thermal denaturation experiments. T_mp_ values are obtained from the native β-sheet peak height measured on the protein in phosphate solution (0.4°C/min heating rate) and in crystalline state (heating rates of 0.4 °C/min and 1 °C/min). Values are calculated by fitting the data with the Boltzmann function. Error bars are the standard deviation from two-four independent experiments. Orange stars indicated T_mp_ values reported in the literature for the three variants in solution (see main text for details). **(C)** Temperature dependence of the tyrosine peak positions (ring ν(CC) mode), taken from the second derivative spectra, of β2m variants in solution and in the crystalline states. The data from the same samples of panel (A) are reported. Orange arrows point to the peak positions reported in the literature (see main text) for deuterated native β2m at neutral pH (orange *) and for deuterated amorphous aggregates obtained after heat treatment of the protein at neutral pH (orange **). The red bar indicates the range of Tyr peak positions here observed for the native soluble variants at room temperature in deuterated crystallization solution.

The loss of native secondary structures upon temperature increase was monitored on crystalline samples at two different heating rates, 0.4 °C/min and 1 °C/min (**Figure** 4). Along the temperature ramps, FTIR signal shifts from native beta structures (component at ∼1628 cm^−1^) to the formation of protein aggregates (bands at ∼1619 and ∼1684 cm^−1^). Under these conditions, at temperature around 90°C the intensity of such component decreases in parallel to the increase in signal corresponding to random coil (band at ∼1648 cm^−1^), indicating a further protein unfolding at high temperatures (**Figure** S3A). This behaviour is evident in the case of the less stable variants: the wt and the D76N β2m (**Figure** 4). In general, at high temperature the protein in the crystallization conditions displays a higher content of random coils and a lower content of intermolecular β-sheets in comparison with sample in PBS (**Figure** 4 and **Figure** S2 and S3).

At the two heating rates similar spectral changes in β2m crystalline samples were detected but differences are evident in *T*_*mp*_ values, calculated from the native β-sheet component ~1628 cm^−1^. Comparable stabilities have been observed for the three protein variants at 0.4°C/min while the T_mp_ for crystals heated at 1°C/min clearly indicated remarkable differences in stability for the W60G and wt β2m compared to the D76N mutant (**Figure** 5). T_mp_ observed at 1 °C/min were found remarkably higher compared to the ones calculated from the experiments at 0.4°C/min (**Figure** 4 and 5). These observations strongly indicate that the stability trends observed for all crystalline samples have relevant kinetic components, likely due to crystal dissolution and to the formation of protein aggregates as indicated by the specific IR marker bands. Interestingly, rapid temperature increase in the experiments at a heating rate of 1°C/min alters the unfolding pathway reducing the formation of protein aggregates. Indeed, in all the investigated samples the structural changes induced by the thermal treatments were found to be irreversible as indicated by the spectra collected after cooling the samples down to room temperature (29), where the FTIR bands typical of protein aggregates are still present at high intensity (**Figure** S2).

The comparison of the T_mp_ calculated from the temperature dependence of the native β-sheet peak height for crystalline and soluble samples indicated higher T_mp_ values for the three variants in the crystalline state (**Figure** 5). The increased stability of the protein crystals is particularly evident in the case of the less stable variants (D76N and wt β2m). This behaviour is in accordance with the downshift of the intramolecular β-structure component of the protein in the crystalline state thus linkable to the more ordered and tight structure of the protein under these conditions.

Importantly, the thermal stability trend for the three variants obtained in solution as well as in the crystalline states at a heating rate of 1 °C/min, is in keeping with the ones previously reported (W60G > WT > D76N) (19, 29, 31, 38) (**Figure** 5).

### Temperature-induced conformational changes monitored by tyrosine FTIR absorption

Further comparative analysis can be performed on the different events occurring during protein denaturation in the solution and crystalline states by monitoring the Tyr ring ν(CC) band at around 1514 cm^−1^ (39). The peak position of this band reflects Tyr chemical environment, therefore providing information on the protein tertiary structures. In the thermal treatments of the β2m variants in solution the Tyr peak positions were found to change by less than 1 cm^−1^, either in deuterated phosphate buffer or in the crystallization conditions (**Figure** 5C). These results agree with the Tyr peak positions previously reported (40) for deuterated native β2m (~1515 cm^−1^) and for deuterated amorphous aggregates obtained after heat treatment of the protein at neutral pH (~1514 cm^−1^).

Conversely, in the crystalline samples the peak position of the Tyr peak is upshifted of more than 2 cm^−1^ during the temperature ramp (**Figure** 5C). The higher shift observed for the crystalline samples (Δ wavenumber > 2cm^−1^) compared to the proteins in solution (Δ wavenumber <1cm^−1^) indicates a relevant change in Tyr environment during the thermal treatments of the crystals, likely reflecting crystal melting. Indeed, the Tyr residues embedded into the crystal “are forced” to interact with the surrounding residues while when crystals melt and release protein molecules in solution, the chemical environment around the Tyr residues undergoes a drastic change. Remarkably, the conformational changes detected by this spectroscopic probe along the thermal treatment of crystalline samples indicate a stability trend in agreement with the measurements of native beta structure in solutions and in crystals: W60G > WT > D76N (**Figure** 5C).

## Discussion

The present work aims at clarifying the extend at which biophysical properties such as protein dynamics and stability assessed *in crystallo* are representative of protein behaviour in solution. Wt β2m and the two mutants D76N and W60G were chosen because of the detailed structural and biophysical characterisation already available. Simulations on wt and D76N β2m result in ensembles which are consistently more rigid when restrained with NMR chemical shifts determined in crystals, as compared to those calculated using solution chemical shifts. The most relevant features emerging from the simulations, however, are preserved in the two environments. Notably, the effects of D76N mutation are qualitatively similar and a comparable scenario can be derived in crystals and in solution. Indeed, all simulations describe an excited state which is particularly flexible for the D76N variant and significantly differs from the native ground state. Conversely, the excited state observed for wt β2m closely resembles the native ground state. Therefore, the systematic comparison of MD simulations performed using solution and solid-state NMR data fully agree with our recent finding that crystals preserve ‘hidden’ conformations reminiscent of pathological states observed in solution (16, 32).

In addition, this new analysis reveals the population of solvent-exposed conformations of D76N β2m W95 side-chain, in both protein states. The position of W95 is central in β2m hydrophobic core and its mutation greatly destabilises the protein fold (21, 34). It is likely that β2m conformations with an exposed W95 may be highly destabilised and may represent an important step in the folding-unfolding pathway. While this conformation could not be detected by X-ray crystallography, it may be a key to understand the low stability of the protein. In contrast, the more stable wt β2m did not show any evidence of such W95 exposure to solvent.

In order to go further in the description of protein crystal properties, protein thermal stability in crystals was then compared to the ones in solution. Melting temperatures of wt, D76N and W60G β2m crystals were in good agreement with previous data in solution (29, 38, 41). This confirms the relevance of assessing fold stability of proteins in crystalline form. Also for this parameter, some systematic differences are observed between crystals and solution samples. T_mp_ values are higher in crystals than in solution, an effect likely due to favourable interactions present in the crystal lattice. Such contribution is more notable for the unstable D76N variant compared to the highly stable W60G (**Figure** 5B). However, the general trend of protein stability is identical in solution and *in crystallo*: W60G mutant is more stable than wt β2m, D76N is the least stable of the three variants. All temperature ramps presented in this work induce irreversible protein unfolding and all the observed processes are non-thermodynamic. Thus, it is not surprising that in the experiments on crystals the kinetic component is relevant and specifically so because crystal melting and protein unfolding appeared as concomitant processes, as additionally suggested by the change of the Tyr peak position. However, at different heating speeds, the T_mp_ values are different but the underlying stability of each of the β2m variants is prevalent and the stability trend is identical in all our experiments. In other words, these data suggest that the experimental setup may change specific values but not general trends in comparative studies adding further to the solidity of this kind of experiments. In summary, the present data suggest that biophysical properties of proteins such as protein dynamics and protein stability are retained in crystals and therefore they can be reliably studied *in crystallo*.

These results are of particular importance to understand and analyze solid-state NMR data. Indeed, solid-state NMR has the unique advantage to probe dynamical processes on different timescales in hydrated crystals or microcrystals. The recent major advances in magic-angle spinning techniques have widened the size range of proteins which can be investigated with site specificity. Here, although we chose to investigate processes - folding and unfolding - implying important conformational changes, the experimental data and simulations indicate that ‘hidden’ conformations of proteins in crystals explain their stability and amyloidogenic properties as consistently as their solution counterpart does. This underlines the pertinence of such approach for the investigation of biological processes where conformational dynamics plays a role.

In addition, the results outline an important role for FTIR to characterise secondary structure content and protein stability on bulk crystalline samples. This technique has proven to be applicable to very diverse kind of samples from cells and tissues to soluble and aggregated proteins (26–29, 35, 37, 42), but, to the best of our knowledge, this work is the first example where FTIR has been used in this context. Crucially, FTIR spectra collected at increasing temperature on crystalline samples were shown to provide specific information on protein unfolding, aggregate formation and crystal melting. This was possible by monitoring the temperature-dependent variation in intensity of the native and inter-molecular beta-sheet components and Tyr peak position.

Previously we showed that FTIR spectra on single crystals (29) provide more insightful information as compared to the spectra recorded in solution, displaying better resolved components. This suggests that crystals are ideal samples to acquire FTIR spectra of excellent quality. However, in our previous work the handling of single crystals was not straightforward and high-quality spectra were successfully collected thanks to the excellent stability of β2m crystals (29). Conversely, crystal pellets are easier to prepare and are more amenable samples, resulting in easily reproducible measurements. Crucially, FTIR spectra collected on bulk crystalline samples provide data of the same excellent quality as single crystals allowing a broader application of this technique in different contests.

Overall, our data indicate that variations of protein stability can be assessed in crystals. This observation suggests that any effect which triggers a variation in protein stability may be appreciated by FTIR experiments: for example, protein stabilisation upon ligand binding may give rise to crystals with an increased stability. Thus, FTIR unfolding experiments may be preliminary or complementary to diffraction data collection to assess the formation of protein complexes. This would be specifically useful in cases of poorly diffracting crystals or limited accessibility to X-ray sources.

Previous reports stressed the relevance of assessing stability of crystals when they are used for industrial application, as materials or as catalysts (43, 44). Crystal stability has been previously assessed using differential calorimetry (43, 44). Our data put forward FTIR as a new insightful and easy-to-use technique to assess crystal stability by monitoring secondary and tertiary structure unfolding and protein aggregation simultaneously.

## Conclusions

In conclusion, here we have presented a detailed comparison of protein dynamics and fold stability using crystalline and solution samples. Our data reinforce the picture that crystals contain a wealth of information about intrinsic properties of proteins such as dynamics and stability and thus, data from crystalline samples may provide deep insights into phenomena such as unfolding and aggregation.

## Author Contributions

BMS, AN, CC and TLM performed the experiments; CC, AN and SR designed the research; BMS, AN, CC, GP TLM and SR analysed the data; BMS, CC, AN and SR wrote the paper with the help of all Authors.

## Acknowledgements

We acknowledge CINECA for an award under the ISCRA initiative, for the availability of high-performance computing resources and support. This work was partly supported by grant of the University of Milano-Bicocca (Fondo di Ateneo 2018-ATE-0284) to AN. This work was partially supported by Fondazione ARISLA (project TDP-43-STRUCT) and by Fondazione Telethon (GGP17036) to SR.

